# microRNA expression during early development in the coral *Acropora digitifera*

**DOI:** 10.64898/2026.05.09.724056

**Authors:** Mila Grinblat, Arie Fridrich, Ira Cooke, Yehu Moran, Roger Hürlimann, Ramona Brunner, Natalia Andrade-Rodriguez, Nobuo Ueda, Eldon E Ball, David J Miller

## Abstract

*Acropora spp*. are the dominant reef-builders of the Indo-Pacific but are also amongst the most stress-sensitive corals. For these reasons, *Acropora spp.* have become the most studied of corals, two species (*A. digitifera* and *A. millepora*) often essentially serving as the basis for understanding molecular responses and processes across the sub-order Refertina and corals in general. The early development of these species has been well-characterised in terms of morphology and gene expression but as yet we have a limited understanding of how transcription is regulated during development. In “higher” animals (bilaterians) microRNAs (miRNAs) are critical regulators of gene expression but until now their involvement in coral development has not been investigated. Building on the existing developmental data for *Acropora spp*., we catalogued microRNAs (miRNAs) expressed during the early development of *Acropora digitifera* and profiled their expression in 21 stages from unfertilised eggs to 24h after treatment with a natural settlement cue (CCA chips). 157 miRNAs were recognised, many of which (∼60%) were novel. These fell into three distinct groups, corresponding to three distinct developmental phases: (1) those present in eggs through to gastrulation (2) a larvally expressed group and (3) those expressed following settlement induction. Exposure of competent larvae to a natural settlement inducer resulted in major changes in the miRNA profile within 10 minutes, indicating that miRNAs may be particularly important in mediating the larva/polyp transition but are also likely to play important regulatory roles throughout early coral development in addition to possible roles in disease resistance.

## INTRODUCTION

MicroRNAs (miRNAs) are small (∼21–23 nt), endogenous RNA molecules that regulate gene expression post-transcriptionally in most multicellular organisms. By binding to complementary mRNA sequences, miRNAs act to repress translation or promote degradation (Bartel 2004, 2009; Moran *et al*. 2014) and play critical roles in regulating development, apoptosis, the cell cycle and many other processes (Pasquinelli et al. 2000; Axtell and Bowman 2008; Horie et al. 2009; Cao et al. 2010; Babenko et al. 2012; Zheng et al. 2012). Bilaterians rely on Drosha and Pasha/DGCR8 for cropping primary-miRNA (pri-miRNA) in the nucleus, followed by cytoplasmic Dicer processing and Argonaute-mediated repression, usually through translational inhibition based on complementarity between the seed-region (nucleotides 2-8 of the guide strand) of the miRNA and the 3’-UTR of its target mRNA (Bartel 2009; Kim *et al*. 2009; Moran *et al*. 2017). In contrast, plant miRNAs undergo both processing steps in the nucleus via Dicer-like 1 (DCL1), HYL1 and Serrate and following their biogenesis, typically cleave mRNA targets after binding via near-perfect complementarity over much longer regions (Cerutti and Casas-Mollano 2006).

In bilaterian animals, the evolutionary turnover of miRNAs is significantly lower than in plants, (Fahlgren *et al*. 2010; Moran *et al*. 2017) leading to >30 miRNA families being conserved across the Bilateria (Prochnik *et al*. 2007; Moran *et al*. 2014). However, despite obvious differences at the functional level, structural similarities of some components imply a common origin for the miRNA processing systems of extant plants and bilaterians, but also with the siRNA processing apparatus (Cerutti and Casas-Mollano 2006; Axtell and Bowman 2008).

Among non-Bilateria, canonical miRNAs are present in cnidarians (Moran *et al*. 2013) and have also been reported in sponges (Robinson *et al*. 2013; Robinson 2015; Liew *et al*. 2016), although the latter appear to be lineage-specific and structurally variable (Calcino *et al*. 2018). No miRNAs have been detected in placozoans and the situation in ctenophores remains unclear (Maxwell *et al*. 2012). There is a clear consensus that cnidarians are the closest outgroup of the Bilateria (for example, Erwin *et al*. 2011; Holzem *et al*. 2024), as reflected in many aspects of their biology. Counterintuitively, however, cnidarians more closely resemble plants not only in terms the biogenesis of miRNAs (see above) but also in that their predominant mode of action involves recognition of mRNA targets via high complementarity over 16-18 bases and cleavage of the transcript within that region (Moran *et al*. 2013, 2014, 2017; Modepalli *et al*. 2018). However, in addition to cleavage of mRNA targets, miRNAs are likely capable of gene silencing via the translational inhibition mechanism that is the predominant regulatory mode in the Bilateria (Mauri *et al*. 2017). Components enabling this process in bilaterians are GW182 proteins, which recruit the CCR4-NOT deadenylation complex via their GW domains, leading to mRNA degradation. GW182 homologs are present in cnidarians (eg. AGW15604 in *N. vectensis*) where they presumably fulfil the same function (Moran *et al*. 2013; Mauri *et al*. 2017).

Of the138 miRNAs known in the anthozoan cnidarian, *Nematostella vectensis* (Praher *et al*. 2021), remarkably miR-100 is the only miRNA shared between cnidarians and bilaterians (Grimson *et al*. 2008) and this has been lost in *Hydra* spp. (Moran *et al*. 2014). Similarly, levels of conservation between both *Hydra* and anthozoans (2 miRNAs) and between the sea anemone *Nematostella vectensis* and corals (9 miRNAs) are also very low (Praher *et al*. 2021). Even within the genus *Hydra*, only 50% conservation was observed between 2 strains (Krishna *et al*. 2013).

Amongst cnidarians, experimental investigation of miRNA function has been undertaken only in the sea anemone, *N. vectensis* (Moran *et al*. 2014; Modepalli *et al*. 2018; Fridrich *et al*. 2020, 2023; Tripathi *et al*. 2022; Admoni *et al*. 2025). Given the depth of the divergence involved (>400 my; Tortorelli *et al*. 2022), conservation of miRNA function between *N, vectensis* and corals is expected to be very low. However, a small number of likely (Fridrich *et al*. 2023) and other potential (Moran *et al*. 2014; Praher *et al*. 2021) exceptions to this general rule have been identified (see below).

To date, possible functions of coral miRNAs have received little attention; the limited literature has focused on resistance/tolerance of *Acropora* spp. to stressors. (Gajigan and Conaco 2017) investigated miRNA expression changes in *Acropora digitifera* during thermal stress. Likewise, Despard *et al*. (2025) investigated miRNA expression levels in *Acropora cervicornis* colonies that were resistant or more susceptible to white band disease. Of the 67 miRNAs identified in the latter study, three were differentially expressed in resistant and susceptible corals (miRNA-33-c, miRNA-2-3p-c and miRNA-2022), the latter two of these being expressed at higher levels after longer exposure to the disease (Despard *et al*. 2025). Single cell analysis of the post-injury responses of *A. muricata* led to the identification of major transcriptomic changes during 14-28d post injury (dpi), whereas miRNA levels were unchanged between the start of the experiment and 28dpi (Han *et al*. 2025).

In bilaterians, as well as roles in cell differentiation, miRNAs are often key regulators of gene expression during early development (e.g. (Rosa and Brivanlou 2017; Divisato et al. 2020), and the (Moran *et al*. 2014) paper on *Nematostella* implies that this is likely to also be the case for cnidarians. To investigate this more extensively, we characterised the miRNA profiles of 15 stages during the early development of *Acropora digitifera*, a common staghorn coral. These data complement our recent transcriptomic analysis of the congeneric species *Acropora millepora* (Brunner *et al*. 2026), providing a broader perspective of the molecular bases of early development of the reef-building corals.

## METHODS

### Sample collection

6 colonies of *A. digitifera* (∼ 25 cm diameter) were collected on 14/6/2019 at the reef in front of Onna, Okinawa, Japan. Corals were brought to the Okinawa Institute of Science and Technology (OIST) Marine Center where they were placed in flow-through aquaria. They were acclimated in filtered sea water (FSW; 1 μm) and a fragment from the middle of each colony was sampled (snap frozen in liquid nitrogen). On the third night after the full moon during the main spawning event in June 2019, colonies were placed into individual containers for oocyte-sperm bundle collection. After separating male from female gametes using a 60 μm mesh, 3 genetically distinct crosses were made by mixing sperm from three separate pairs of colonies (oocytes from one member of the pair, and sperm from the other) and raised in separate tanks for the duration of the experiment. Samples of adult colonies, unfertilized oocytes and 12 developmental stages (four cell stage to competent larva) were collected as listed in Tables S1 and S2. Prior to sampling, the developmental stage of each cross was confirmed based on morphological characteristics; for reference, SEM images showing the early development of a typical *Acropora* spp. are provided in Figure 1 of Brunner *et al*. (2026). That paper also introduces vernacular terms for early developmental stages (e.g. “prawn chip”). The motile (ciliated) larval stage that many cnidarians go through is often referred to as the planula and in this manuscript, “larva” and “planula” are used interchangeably.

**Figure 1.**
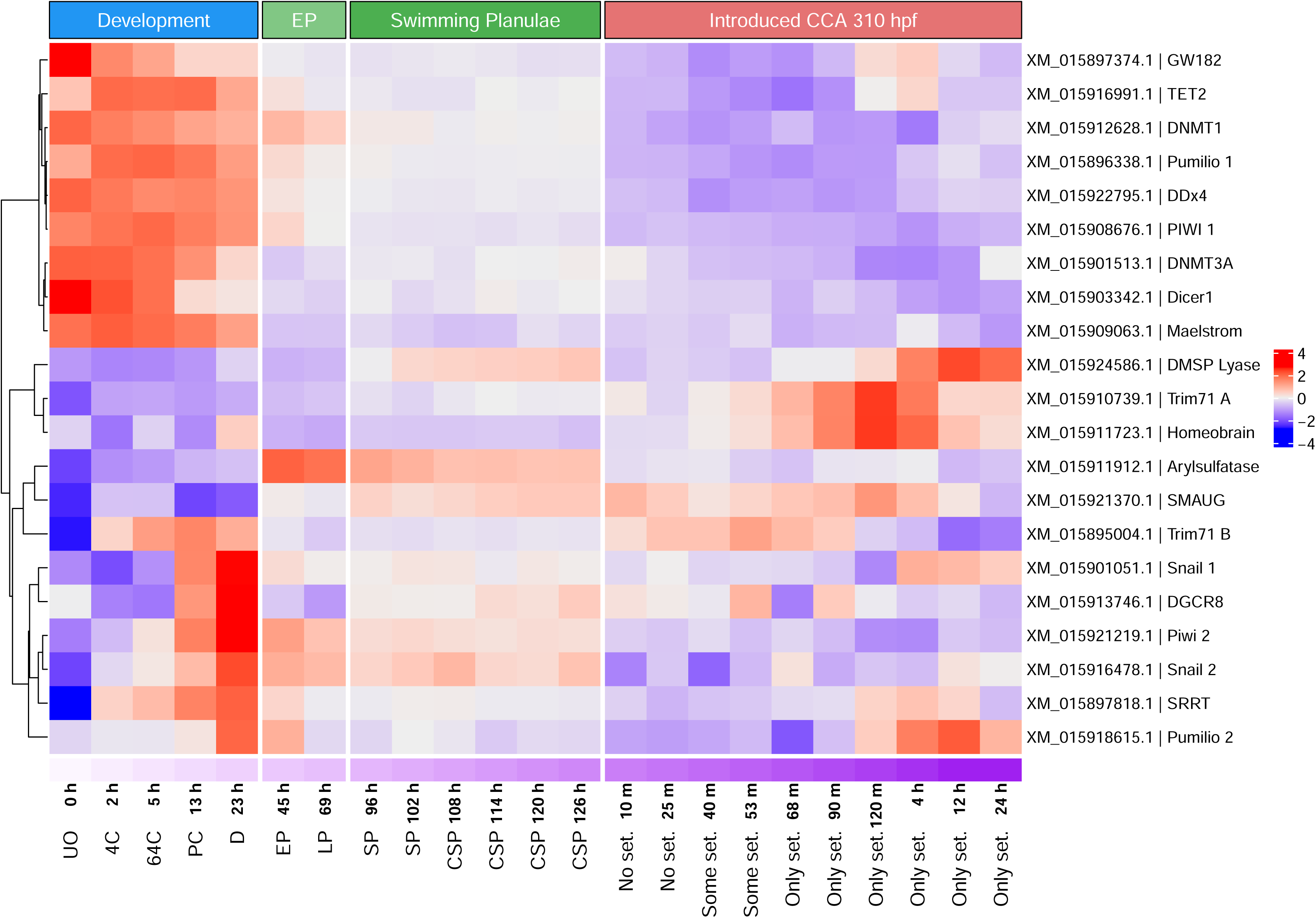
Expression of specific marker genes during the early development of *Acropora digitifera*. Columns are ordered by stage/time point, from 0 to 126h post fertilisation, followed by 10m to 24h after introduction of the settlement cue (CCA; at 310hpf). Rows show RefSeq transcript IDs and gene names for marker gene transcripts (see Table S3), including developmental regulators, components of the miRNA-processing and RNA-silencing machinery, and putative miRNA target genes. Stages are abbreviated across the bottom of the figure as: unfertilised oocyte (UO), 4-cell (4C), 64-cell (64C), prawn chip (PC), doughnut (D), early planula (EP), late planula (LP), swimming planula (SP), and planula competent for settlement (CSP); developmental stages for which vernacular terms have been applied are illustrated in Figure 1 of Brunner et al. (2026). Post-settlement expression levels are labelled by time after introduction of the settlement cue (CCA chips). Colours indicate relative expression levels and represent the means of the three biological replicates.

### Settlement

Around 310h post-fertilisation, approximately equal numbers of larvae from each of the crosses (A, B and C) were pooled in an additional tank, to be used for miRNA analyses. Whilst miRNA libraries were prepared only from pooled larvae, samples from each of the three crosses were used for RNASeq (mRNA) analyses at time points matching those for miRNA analyses. In both cases, aliquots of 100 larvae from each cross were placed in separate batches of petri dishes (10 per cross) each containing 20 ml of FSW. This was repeated for each time point (10 time points, total 40 petri dishes). At time zero, 5x5 mm chips of crustose coralline algae (CCA) were introduced to each of the dishes and larvae were sampled from a designated dish at each time point. 90 min after introducing CCA, only larvae that settled were used. All samples were fixed in RNA Shield (Zymo) and stored in a freezer at −20C.

### RNA extraction and sequencing

Total RNA from adult coral fragments was extracted using Trizol (Thermo Fisher) and cleaned using the RNA Clean & Concentrator Kit (Zymo, R1017) following the manufacturer’s instructions. Total RNA from development and settlement stages was extracted using the miRNeasy Mini Kit (Qiagen) following the manufacturer’s instructions using TissueLyser LT (Qiagen) as homogenizer. Quality of RNA was assessed using 4200 TapeStation (Agilent) and only samples with RINe number over 7 were used.

Small RNA libraries were prepared using the Small RNA-Seq Kit v3 for Illumina® Platforms (NEXTFLEX®) and sequenced on one lane of an Illumina Hiseq2500 resulting in between 8 and 19 million 50 bp paired end reads per sample. RNA-seq libraries were prepared using NEBNext Ultra II Directional RNA kit for Illumina and sequenced on a single S1 flow cell on a NovaSeq6000 resulting in between 19 and 50 million 150 bp paired-end reads per sample. Library preparation and sequencing of both miRNA and mRNA were performed at the OIST sequencing facility. Samples used for miRNA and mRNA analyses are listed in Tables S1 and S2, respectively.

### Post sequencing analyses- miRNA

The raw reads were processed according using a workflow based on (Mohammed and Hekmaty 2018). Briefly, 3’ adapter sequence was clipped (TGGAATTCTCGGGTGCCAAGG) then the 4 first and 4 last bases were trimmed from the adapter-clipped reads. Remaining reads were mapped to the *A. digitifera* genome (GCF_000222465.1_Adig_1.1_genomic.fna) using Bowtie (v.0.12.7) and mapper.pl as part of the miRDeep2 ver. 2.0.0.8 package (Friedländer *et al*. 2012) with default parameters discarding reads smaller than 18bp. The total number of miRNA reads processed averaged 27,708,601 per sample with an average of 14,178,036 (∼51%) mapping and 13,530,565 (∼49%) failing to map to the genome (Table S1). Novel and known miRNAs were identified using miRDeep2.pl core algorithm using a list of miRNAs known from other cnidarian species as input (Table S3, (Krishna et al. 2013; Liew et al. 2014; Moran et al. 2014; Gajigan and Conaco 2017; Baumgarten et al. 2018; Urbarova et al. 2018; Fridrich et al. 2020; Praher et al. 2021; Despard et al. 2025). Additionally, provisional mature miRNA sequences were matched to the known mature miRNAs in Table S3 by direct sequence comparison using custom code implemented in R (R Core Team), retaining only best matches with ≥90% sequence identity. Any novel potential miRNA was considered valid only if it passed the criteria previously published (Fromm *et al*. 2015; Fridrich *et al*. 2020). The quantifier.pl module (miRDeep2 package) with default parameters was used to quantify miRNA expression. The counts were normalized to library size using edgeR (Robinson *et al*. 2010; McCarthy *et al*. 2012). DE analysis was performed using the limma Package (Ritchie *et al*. 2015). For DE analysis “tank” was treated as a random effect using the duplicateCorrelation function in limma. Heat maps were generated using ComplexHeatmap (Gu *et al*. 2016). Rows were partitioned into three clusters using k-means clustering (row_km = 3) to distinguish between miRNAs that are upregulated in early development, planula and settlement stages, and rows within clusters were ordered by hierarchical clustering using Pearson correlation distance (clustering_distance_rows = “pearson”). The cluster names were assigned according to the main expression pattern; HI_DEV, included miRNAs that were mostly highly expressed during development, HI_PLAN, during planula stages and HI_SET, during settlement. An MDS plot was generated using the plotMDS function from edgeR.

As the expression pattern of sample AD_45 (miRNA, collected 00:53 min after CCA was introduced) was significantly different from the rest of the settlement samples (MDS) possibly due to a technical issue with the sample processing, it was removed from the miRNA analyses. The original MDS plot is available in Figure S1.

### Post sequencing analyses - mRNA

Adapters were trimmed using trimgalore v0.6.11 (Martin 2011). STAR Aligner v2.7.11b (Dobin *et al*. 2013) was used to map the reads to RefSeq transcripts for *A. digitifera* (GCF_000222465.1) and to generate gene counts for each transcript. Quality control was performed using rseqc v5.0.4 (Wang *et al*. 2012) and multiqc v1.33 (Ewels *et al*. 2016). DE analysis with contrast was performed using the Limma Package v3.66.0. For the DE analysis “tank” was treated as a random effect. Heat maps of all differentially expressed mRNAs as well as genes of interest were generated using ComplexHeatmap. The MDS plot was generated using the plotMDS function from edgeR.

### miRNA target prediction

Targets were predicted using psRNATarget v2 (Dai and Zhao 2011; Dai et al. 2011, 2018), with expectation maximum of 2.5 allowing no gaps as previously used in Fridrich et al. (2020). Targets were annotated as described in https://github.com/iracooke/acropora_digitifera.

### miRNA-mRNA correlation

miRNAs regulate their targets in different ways, these being known as coherent or incoherent regulation (Shkumatava et al. 2009; Ebert and Sharp 2012). In coherent regulation, miRNA mediated repression reinforces the overall action of the upstream regulatory machinery on the target. In this case the miRNA and the target should show opposing expression patterns (negative correlation). Incoherent regulation occurs when miRNA and the targets are activated by the same regulators and the miRNA simply acts as a fine-tuning mechanism maintaining the precision of the expression of their targets as they buffer stochastic fluctuations in gene expression. In this case the expression of both miRNA and target will, to an extent, overlap temporally or spatially (Shkumatava et al. 2009). Correlation analyses were performed to determine whether miRNA-target regulation was coherent or incoherent. The correlation analysis was performed twice. First, only pre-settlement stages were included, because these stages had directly comparable miRNA and mRNA samples in both datasets (Tables S1 and S2). Second, a separate correlation analysis was performed in which post-settlement stages were included. For this latter analysis, the three mRNA post-settlement samples at each time point were averaged to match the single pooled miRNA post-settlement sample available for the corresponding time point. The 00:53 hpi settlement sample was excluded because it was removed from the miRNA analysis, and the 04:00 hpi and adult coral samples were excluded because they were not present in both datasets. Averaging the mRNA settlement samples was justified because the pooled miRNA settlement samples contained larvae from all three crosses, whereas these crosses were sequenced separately in the mRNA analysis. In both correlation analyses, the mRNA count table was filtered to retain only genes predicted to be miRNA targets. Pearson correlation of miRNAs’ and the targets’ normalized counts were performed using row.pearson function from the HybridMTest package (Release 3.22) (Pounds and Fofana 2025) with p-value < 0.05, correlation values under −0.5 (negative correlation) and over 0.5 (positive correlation) as a cut-off. Heat maps of coefficients were generated using Complexheatmap.

## RESULTS

### The developmental patterns of A. digitifera and A. millepora are similar

The early development of *A. digitifera* closely mirrors that of the congeneric coral, *A. millepora*, in terms of both morphology and gene expression profiles (Ball *et al*. 2002; Grasso *et al*. 2008, 2011; Meyer *et al*. 2011) and the latter has been the subject of a recent comprehensive analysis of early development (Brunner *et al*.. 2026). To complement the *A. digitifera* miRNA analyses, expression levels of a number of genes considered to represent key events in early development (Brunner *et al*. 2026) were determined in some of the same material (see Figure 1). Note, however, that some of the expression profiles did not correspond directly between the two species in timing, which is due in part to differences in sampling intervals but perhaps also to development under different environmental conditions.

One factor that might be expected to have significantly impacted post-settlement development is that, in the case of *A. millepora*, settlement was induced at around 40h after competency was acquired (Brunner *et al*. 2026), whereas in the *A. digitifera* experiment, the period between competency and settlement induction was much longer (184h). The latter scenario may, however, more closely resemble the normal (“real world”) situation. In nature, high proportions of the larvae of many *Acropora* spp. remain competent for >20 days (Connolly and Baird 2010) and delayed settlement had only very limited impacts on growth rates (as measured by time to initial budding) and survival (Graham *et al*. 2013). However, we cannot discount possible effects of time post-competence on the earliest molecular events occurring post-settlement.

The use of somewhat different settlement cues may also have had some impact on early post-settlement development rates. The settlement cue used in the Australian *A. millepora* study was an ethanolic extract of the CCA *Porolithon onkodes* (Brunner *et al*. 2026). In the present case, in Japan, this same extract had little or no impact on competent *A. digitifera* larvae. By contrast, whole chips of local CCA effectively initiated settlement and metamorphosis of *A. digitifera* larvae. Note that such differences between *Acropora* spp. in responses to settlement cues have previously been documented (Abdul Wahab *et al*. 2023).

Nevertheless, the marker gene profiles (Figure 1) were generally consistent with the idea that both the pre- and post-settlement developmental patterns previously documented for *A. millepora* applied in *A. digitifera*; for example, the expression of XM_015901051.1, the *A. digitifera* homolog of snail, at around 13-23h post-fertilisation matches that reported in *A. millepora* (Hayward *et al*. 2004; Ball *et al*. 2021; Brunner *et al*. 2026). The *A. digitifera* Piwi2 (see Praher *et al*. 2017) homolog XM_015921219.1 (expression of which peaks at 23hpf) corresponds to that of *A. millepora* XP_044168784.1; expression of both Piwi2 and Snail homologs marks the first major wave of zygotic gene expression (Brunner *et al*. 2026) and therefore the maternal/zygotic transition (MZT). As in *A. millepora*, changes in the expression of key developmental genes, including homologs of homeobrain and Trim71/Lin41, occurred several hours after settlement induction in *A.digitifera*. Likewise, expression of XM_015918615.1, a homolog of mammalian Pumilio 3, peaked at both early (23hpf) and later (4-12h post-induction) time points in *A. digitifera*, corresponding to the 29hpf and 6-12hpi time windows for XP_044164192.1 (=XM_044308257.1) in *A. millepora* (Brunner *et al*. 2026). On the basis of these gene expression patterns, we assume that *A. digitifera* undergoes a developmental event resembling the “juvenile/adult” transition of *Caenorhabditis elegans,* several hours after settlement induction as was documented in Brunner *et al*. 2026. This event presumably corresponds to the “point of no return” documented by Ishii et al. (2022), which marks the time at which metamorphosis becomes irreversible.

### The miRNA repertoire of *A. digitifera*

A total of 157 miRNA predictions were generated in this study, corresponding to 134 unique mature miRNA sequences after collapsing duplicated predictions that produce identical consensus mature sequences from different genomic loci (Table S4). Of these, 63 correspond to miRNAs previously described in cnidarians, while 94 (85 unique) are newly identified in this study. Among the previously described miRNAs recovered here, 43 have been reported from *Acropora digitifera*. The number of miRNAs identified to date in *A. digitifera* is therefore in the same range as in *Nematostella vectensis* (166) and *Hydra magnipapillata* (126) ; somewhat smaller numbers have been identified so far in several other anthozoans: *Acropora cervicornis* (63; Despard *et al*. 2025), *Acropora millepora* (40; Praher *et al*. 2021), *Stylophora pistillata* (31;Liew et al. 2014), *Anemonia viridis* (83; Urbarova *et al*. 2018) and *Exaiptasia pallida* (51; Baumgarten *et al*. 2018). The smaller numbers of miRNAs identified to date in some species are likely to be underestimates given that they have been based on surveying fewer life cycle stages. Figure 2 summarises the extent of overlap between the known miRNA profiles of the three *Acropora* species that have been investigated so far.

**Figure 2.**
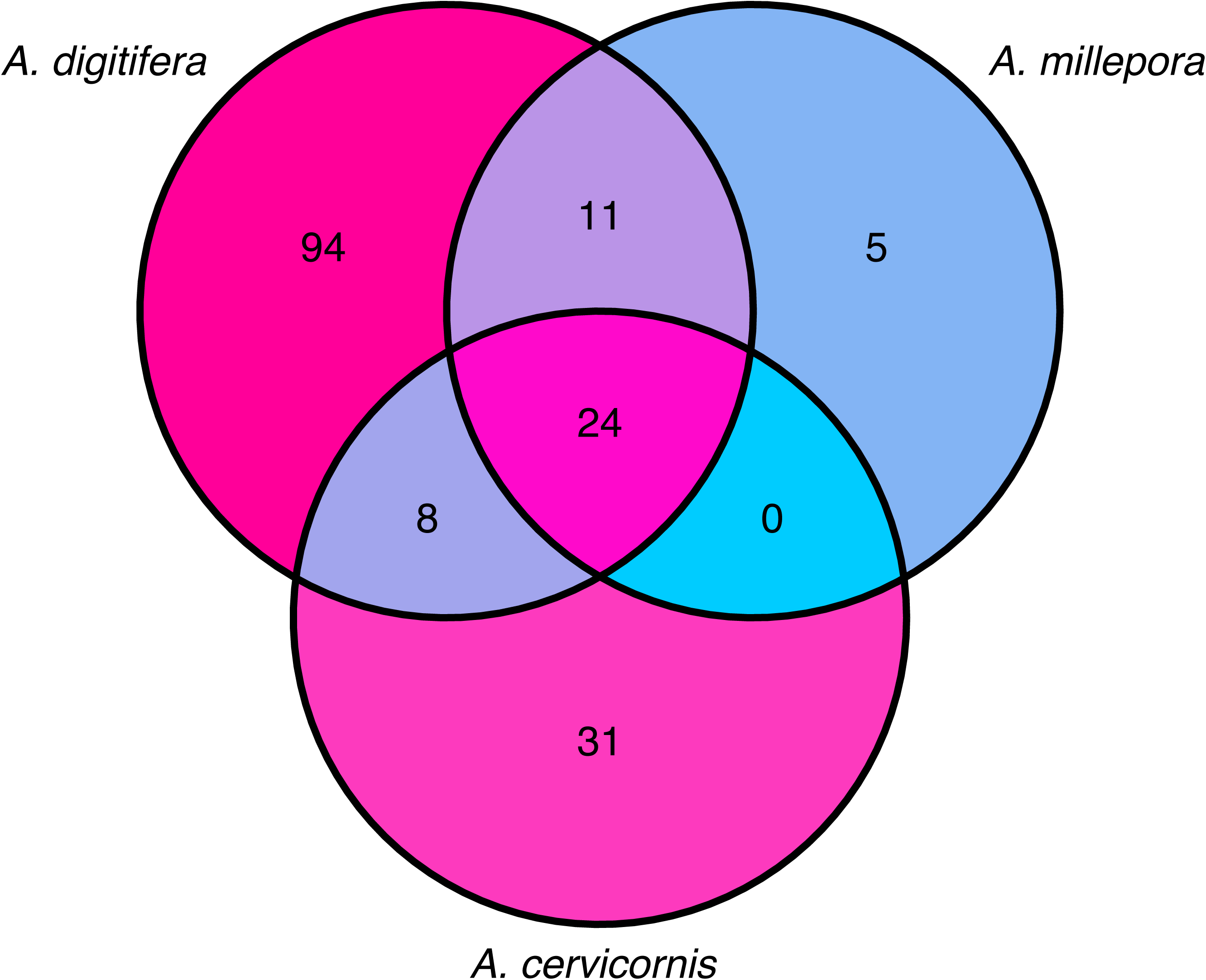
Venn diagram showing the overlap of miRNA IDs among three *Acropora* species (*Acropora digitifera*, *Acropora millepora* and *Acropora cervicornis*). Numbers within each region indicate the count of miRNA IDs unique to, or shared among, the indicated species. miRNA IDs shared by all three species are 1, 2, 3, 4, 5, 7, 8, 10, 13, 14, 17, 19, 24, 29, 33, 100, 2022, 2023, 2025, 2030, 2036, 2037, 2050 and 9425.

Seven of the 9 miRNAs previously reported as shared between *A. digitifera* and *N. vectensis* were identified in the present study (miR-2037 and miR-2050 were not detected).

Comparisons of miRNA conservation are complicated by the *ad hoc* non-miRbase miRNA nomenclature adopted by some authors, which can result in both duplicated identifiers for identical mature sequences and reuse of the same number for unrelated miRNAs. These cases demonstrate that miRNA numbering alone does not reliably reflect homology or family membership. Consequently, estimates of miRNA conservation based solely on numeric identifiers should be interpreted with caution, and sequence-level validation is required to establish evolutionary conservation.

### Differential miRNA expression during development

As a step towards understanding the roles of miRNAs during development in *A. digitifera*, the expression of miRNAs was profiled from unfertilised eggs to primary polyps via Multi Dimensional Scaling (MDS) analysis (Figure 3). A recent study from our group generated a comprehensive gene expression profile of early development from unfertilised egg to primary polyp in the related species *A. millepora* (Brunner *et al*. 2026) which served as a template for the present study. Note that, in the present case, sampling intervals and timing differed somewhat, to account for regulatory changes imposed by miRNAs that might take place prior to changes in gene expression. For the developmental analyses, miRNA data for *A. digitifera* were obtained for 21 developmental stages (Table S1) and a representative adult. In parallel, mRNA analyses were conducted on some of the same material (Figure 1; Table S2), enabling miRNA/mRNA correlation analyses.

**Figure 3.**
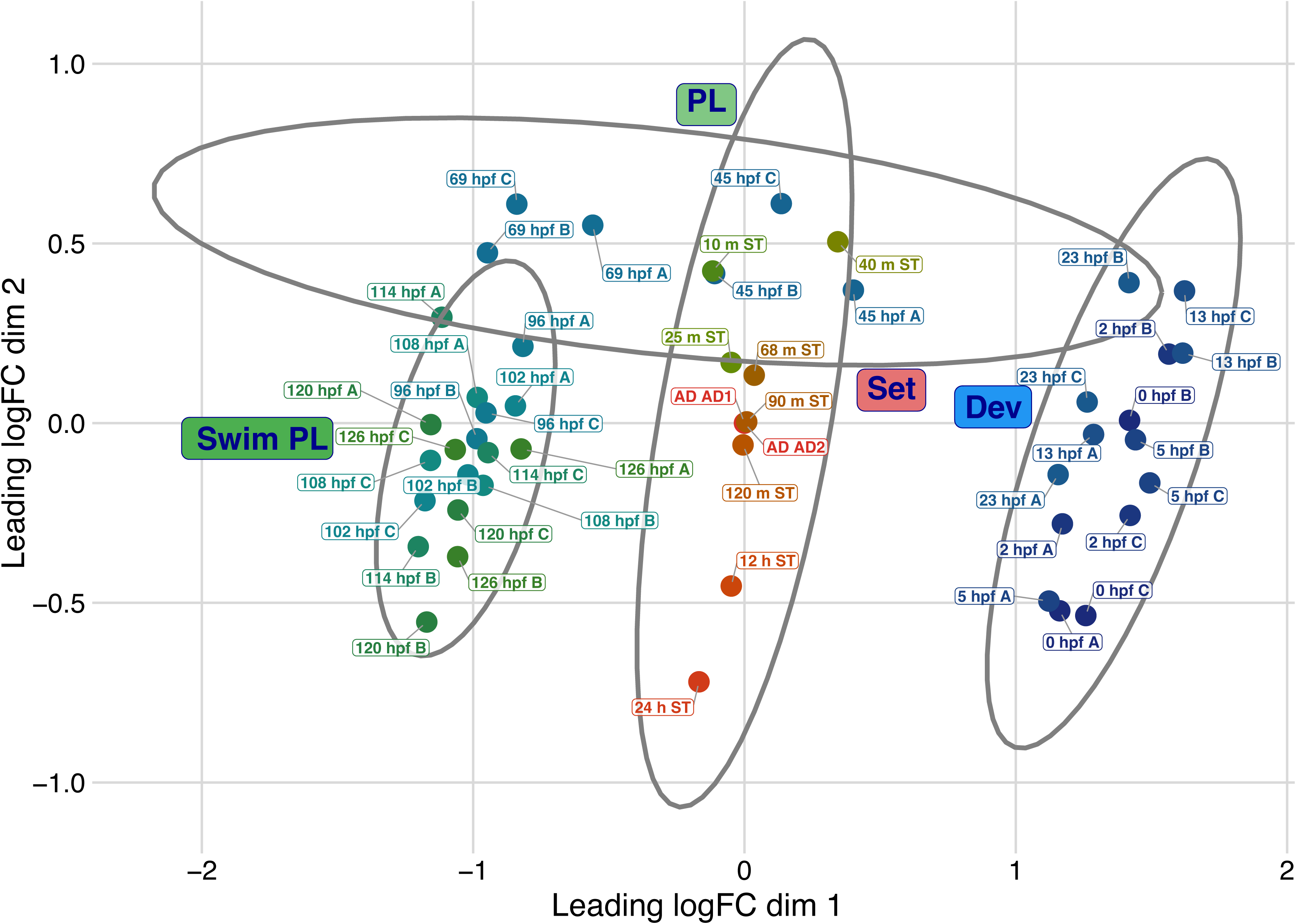
Multidimensional scaling (MDS) ordination of expression profiles across development (Dev), planula (PL), Swimming planula (Swim PL), settlement (Set), and adult samples. The panel shows the leading logFC dimensions 1 and 2. A-C labels indicate sample provenance (tanks A–C, adult colonies AD1–AD2, and the settlement experiment ST), as shown in the key. Settlement samples are labelled by time after introduction of the settlement cue (CCA chips). Grey ellipses highlight the broad groupings labelled Dev, PL, Swim PL, and Set in the figure.

Of the miRNAs that passed the low expression threshold, all 131 were significantly differentially expressed across developmental and settlement stages and with the adult sample (F-test, Figure 4); the 23 miRNAs that did not pass the threshold are listed in Supplementary Table S5. As in the case of the MDS analysis (Figure 3), the differential expression analysis (Figure 4) showed a distinct shift in miRNA expression patterns during the early planula (EP) stage, and another major shift after the introduction of the settlement cue (CCA chips). The adult miRNA expression pattern also differed substantially from that of settling planulae (Figure 4).

**Figure 4.**
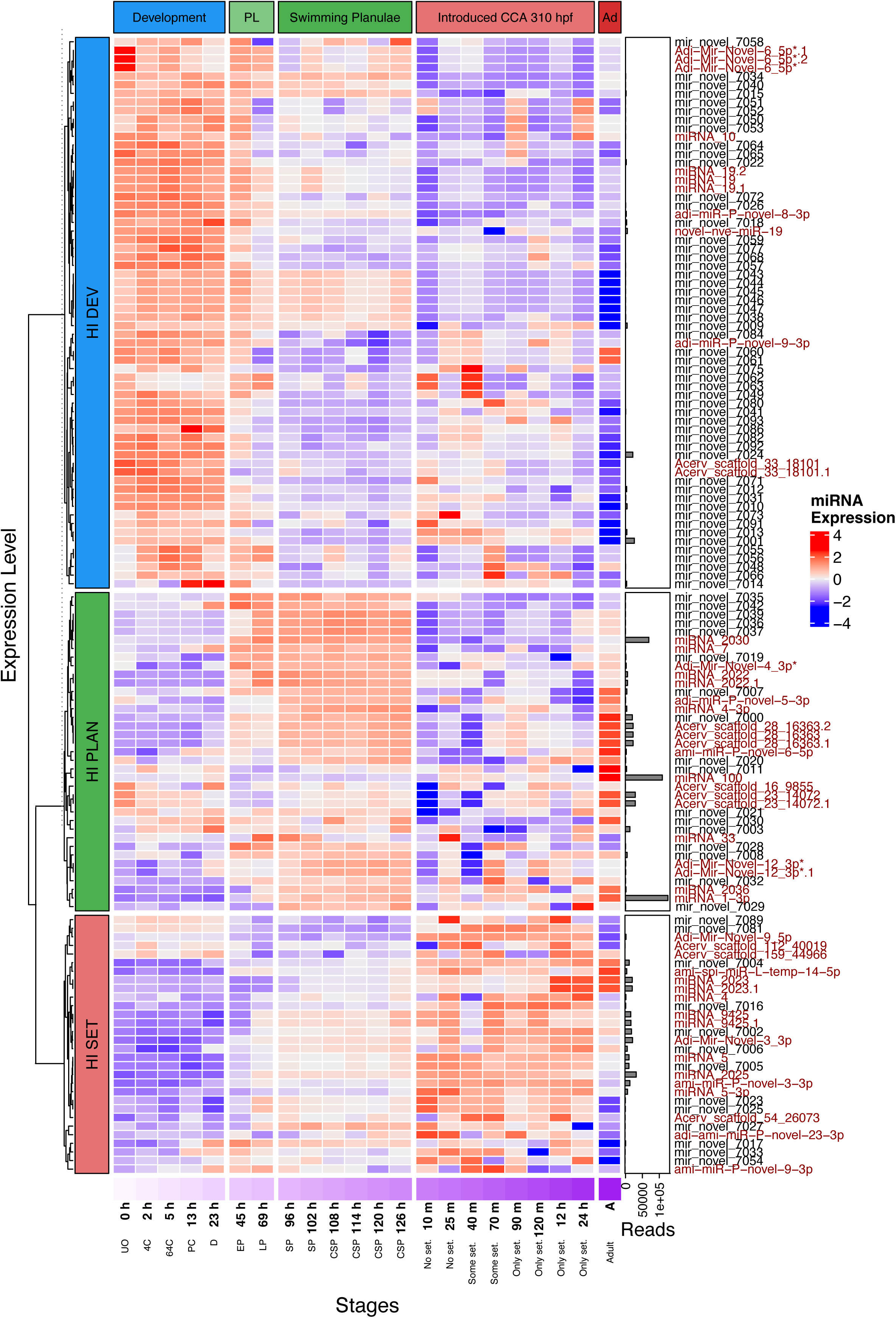
Differential expression of miRNAs through the early development of *Acropora digitifera*. Rows are miRNAs and columns are samples ordered by stage/time point. Relative expression is shown by the colour scale (range shown in the key). Abbreviations for developmental stages shown across the bottom of the figure are as listed in the legend to Figure 1, with the addition of data for an adult individual (AD). Post-settlement samples are labelled by time after introduction of the settlement cue (CCA chips). K-means clustering grouped miRNAs into three expression clusters labelled HI_DEV (predominantly higher during development), HI_PLAN (predominantly higher during planula stages), and HI_SET (predominantly higher during settlement). A bar plot of read counts per miRNA is shown alongside the heatmap. Red text is used to indicate miRNAs that have previously been described.

The application of k-means clustering classified most of the miRNA expression patterns into three groups (Figure 4): (1) cluster HI DEV, where many miRNAs were at their highest levels during development, (2) cluster HI PLAN, where miRNAs were upregulated following gastrulation and expressed throughout the planula phase and (3) cluster HI SET, with most miRNAs upregulated immediately (i.e. within 10m) following exposure to the settlement cue. None of the novel miRNAs was unique to the adult stage, most of those upregulated being detected in the HI PLAN and HI SET clusters, but many of these were at their highest levels in the adult sample.

### Most of the HI DEV miRNAs are maternal in origin

mRNAs encoding some miRNA biogenesis components (Moran *et al*. 2017) in cnidarians are provided maternally in *Acropora millepora*, (e.g. Dicer) whereas others (eg. Serrate/SRRT) are early zygotic genes (Brunner *et al*. 2026). As far as can be determined, the pattern of expression of small RNA processing activities established in *A. millepora* holds true for *A. digitifera* (Figure 1; Supplementary Table S6), implying that most of the miRNAs involved in the first of the three most obvious groups of miRNAs (“HI DEV” cluster) in Figure 4 are maternal rather than zygotic in origin. One clear exception to this pattern is the novel miR-7014 which was first detected at the 13hpf time point, at which embryos are entering gastrulation (i.e. they are at the “prawn-chip” stage).

During the development of *A. millepora*, the mRNA expression profile indicates that, by the early planula stage the full miRNA processing machinery is likely to be in place (Brunner *et al*. 2026), and the expression profiles of homologs of some of these genes confirmed that this is also the case in *A. digitifera* (Figure 1 and Table S6).

From 23hpf, the *A. digitifera* maternal (“HI-DEV”) miRNA profile begins to shift such that by 96hpf (the early swimming planula stage) a second miRNA profile (“HI-PLAN”) is established (Figure 4). The implication here being that the second major cluster of miRNAs (the “HI-PLAN” group) visible in the 45-69hpf time window in Figure 4, includes many of the earliest miRNAs of zygotic origin.

### The HI-PLAN and HI-SET miRNA profiles

miRNA levels within this second cluster appeared to be relatively stable across the late pear to mature planula (i.e. 69 -126hpf) stages (Figure 4). This was unexpected, as this time window includes the transition to competence, suggesting that miRNAs have little involvement in this process in *A. digitifera*.

However, the induction of settlement of competent planulae at around 126hpf immediately (i.e. within 10 minutes) resulted in extensive changes to the miRNA profile (Figure 3). Note that many of these changes in miRNA levels occurred several hours before major changes were observed in the expression of several genes thought to be involved in the larva/polyp transition (Brunner *et al*. 2026). Whilst many of the miRNAs up-regulated on induction formed a distinct cluster (labelled “HI-SET” in Figure 4), overall, the late phase miRNA profile was considerably more dynamic than were the early developmental (“HI-DEV”) or planula (“HI-PLAN”) phases.

The adult miRNA expression pattern was clearly distinct from those of previous stages, some miRNAs (e.g. miR-100-5p) being unique to the adult stage, and a significant number of others being present at their highest levels in adults (Figure 4). Amongst these was miRNA-1-3p; note that this is a classic example of confusing *ad hoc* miRNA nomenclature (as mentioned above). The *Acropora* miRNA miRNA-1-3p reported by Despard et al. (2025) differs substantially from the entry miRBase MIMAT0000416.

### Prediction of miRNA targets and correlation of miRNA/target expression patterns

Sequence complementarity provides a direct approach to the identification of miRNA targets. Although the extensive complementarity characteristic of miRNA/mRNA cleavage interactions means that this approach is more applicable to cnidarians than to bilaterians, false positives remain a major issue (Fridrich *et al*. 2019).Hence this approach is most useful in the context of evaluating whether interactions demonstrated in one species could occur in others.

Application of the psRNATarget software to the *A. digitifera* data predicted a total of 1163 miRNA/mRNA interactions for 121 of the of 157 miRNAs identified, but this number included 131 corresponding to ncRNAs (Table S7).

Analyses of the (negative or positive) correlation of miRNA and mRNA data for putative miRNA targets can reveal the type of regulation by indicating how miRNAs might be influencing their targets (Enerly *et al*. 2011; Allantaz *et al*. 2012). In the case of cnidarians, some miRNAs are co-expressed with their mRNA targets (Moran *et al*. 2014), justifying the application of this approach to the *A. digitifera* data.

Given that miRNA analyses (but not the mRNA analyses) post-induction were conducted on pooled larvae (see methods section) whereas those on pre-induction material were based on three biological replicates (as were the mRNA analyses), the initial round of miRNA/mRNA correlation analyses was conducted based only on the pre-induction data. In these analyses, a total of 27 miRNA-mRNA pairs (excluding ncRNAs) were found to be negatively correlated (Table S8, Figure 5) and a total of 51 pairs found to positively correlate in all stages (Table S9, Figure 6). For completeness, a second round of correlation analyses was conducted, based on the full range of time points. These results are summarised in Tables S10 and S11, and in Figures S2 and S3.

**Figure 5.**
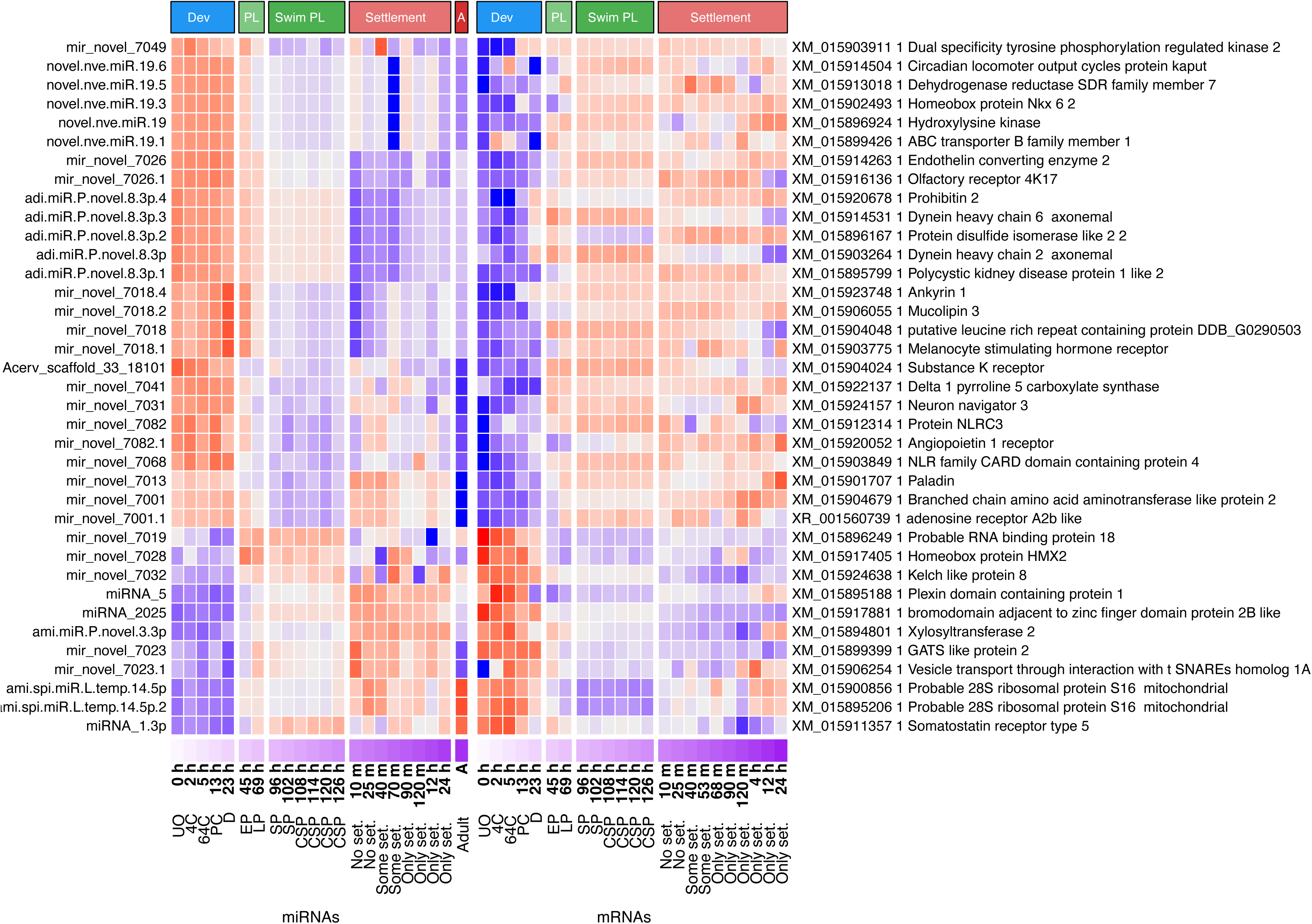
Heatmaps showing expression patterns for miRNA–mRNA pairs with negative correlation (correlation coefficient ≤ −0.5; Table 5). The left panel shows miRNA expression and that on the right shows expression of the corresponding predicted target mRNAs, aligned by sample order. Note that, whereas negative correlation was demonstrated only for the pre-settlement stages, the heatmaps show expression levels across all time points sampled. Paired heatmaps illustrating data for genes showing negative miRNA/mRNA correlation across all time points are shown as Figure S2. Abbreviations used for developmental stages are as in the legend to Figure 1. Post-settlement samples are labelled by time after introduction of the settlement cue (CCA chips). Only targets with functional annotation are shown.

**Figure 6.**
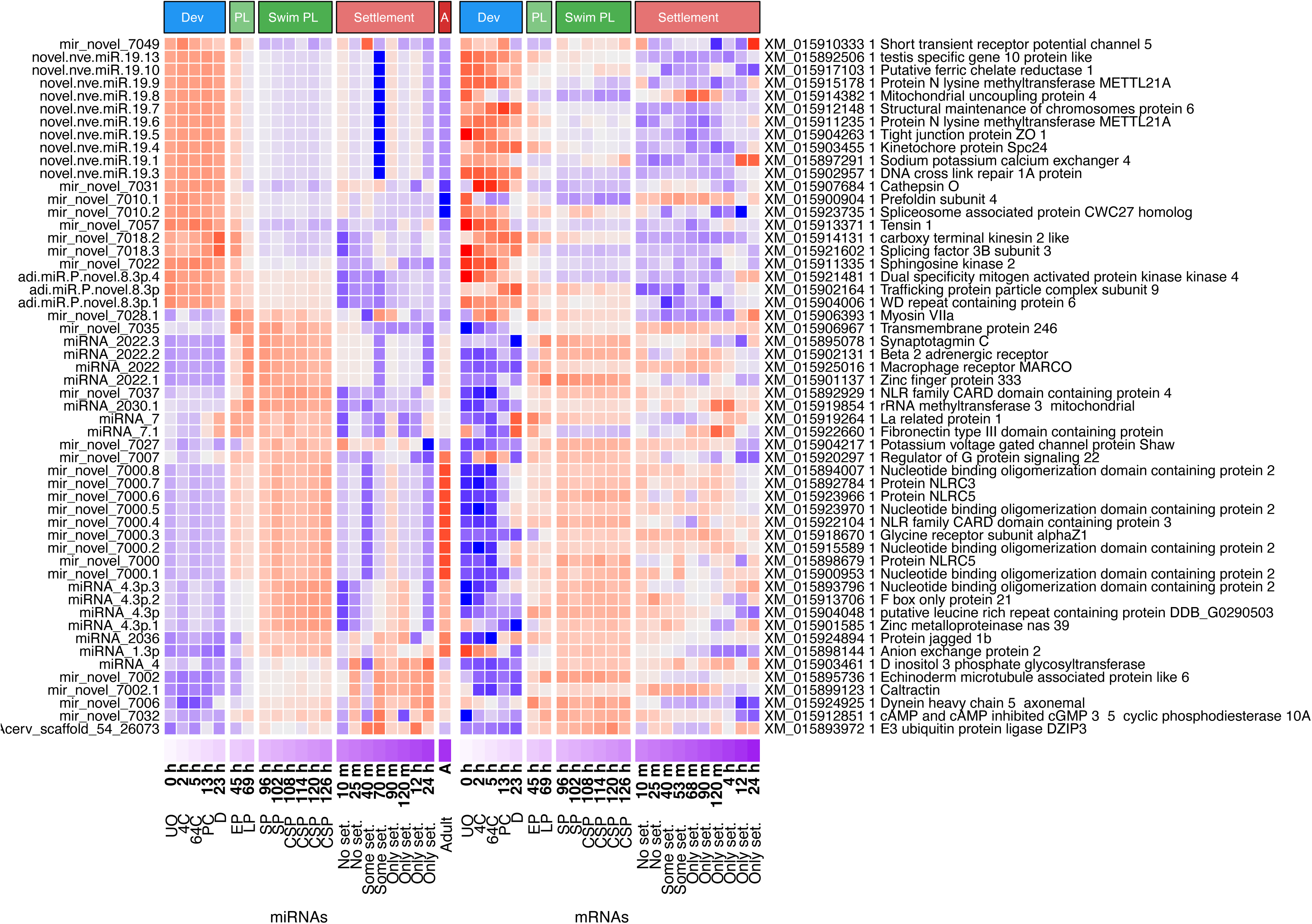
Heatmaps showing expression patterns for miRNA–mRNA pairs with positive correlation (correlation coefficient ≥ 0.5; Table 6). The left panel shows miRNA expression and that on the right shows expression of the corresponding predicted target mRNAs, aligned by sample order. Note that, whereas positive correlation was demonstrated only for the pre-settlement stages, the heatmaps show expression levels across all time points sampled. Paired heatmaps illustrating data for genes showing positive miRNA/mRNA correlation across all time points are shown as Figure S3. Abbreviations used for developmental stages are as in the legend to Figure 1. Post-settlement samples are labelled by time after introduction of the settlement cue (CCA). Only targets with functional annotation are shown.

Note that the lack of obvious correlation does not rule out the possibility of miRNA/mRNA interactions occurring in vivo.

## DISCUSSION

In total, 157 miRNAs were identified in *A. digitifera*, 43 matching previously reported *A. digitifera* miRNAs, 63 had previously been detected in other cnidarians and 40 of these were conserved with the congeneric species *A. millepora* (Praher *et al*. 2021; Table S4). The limited conservation of the miRNA repertoire observed here is consistent with previous work on cnidarians (Moran *et al*. 2014; Fridrich *et al*. 2020; Praher *et al*. 2021) and contrasts markedly with the high level of conservation seen across the Bilateria. Although a higher proportion (52 of 64) of the *A. digitifera* miRNAs in the HI-DEV cluster were novel relative to the HI PLAN (17 of 37) or HI SET (13 of 30) clusters (Figure 4) the most likely explanation for this is that early stages of *A. millepora* (the congeneric species contributing most of the matches with *A. digitifera* miRNAs) were underrepresented in previous (Praher *et al*. 2021; Despard *et al*. 2025) studies.

Note that in addition to direct action via mRNA cleavage, silencing via translational repression may also occur during development in *A. digitifera*. The key components enabling this are members of the GW182 family. Note that XP_015752860.1 (=XM_015897374.1) is the likely *A. digitifera* GW182 homolog, and the fact that the corresponding mRNA is maternal (Figure 1) implies that translational repression may be imposed by miRNAs throughout early development in *A. digitifera*.

### Potential physiological implications of miRNA expression data

Several putative targets of miRNAs are associated with larval motility. As planula larvae of *A. digitifera* and other corals develop, ciliary movement marks a key transition, as in many marine larvae (Staver and Strathmann 2002). Four axonemal dyneins essential for ciliary motion in mice, zebrafish, and *Drosophila* (Olbrich *et al*. 2002; Yamaguchi *et al*. 2018; Li *et al*. 2021), DNAH2, DNAH5, DNAH6, and DNAH8 were predicted targets of *A.digitifera* miRNAs (Table S7). During early development and in the larval stages, levels of mir_novel_7006 and its predicted target DNAH5 were positively correlated (Figure 6; Table S9), implying incoherent regulation. miR-mir_novel_7006 is weakly expressed in swimming planulae, drops briefly when the settlement cue is applied, and then rises again. This profile is compatible with cilia having roles in motile larvae and in post-settlement/adult stages, but not during metamorphosis itself.

Given that pre-settlement development of *Acropora* spp. and many other corals relies on metabolism of storage lipids, the potential interaction of miR-7054 with lipase XP_015769681.1 (=XM_015914195.1 Table S7) may also have developmental significance.

Also of potential physiological significance is the predicted interaction between miR-2022 and DMSP lyase XM_015924586.1. DMSP is a potent antioxidant and is present at high levels in reef-building corals, particularly *Acropora* species. Corals are unique amongst metazoans in their ability to synthesise DMSP (Raina *et al*. 2013), although DMSP synthesis in mature corals is largely brought about by their resident Symbiodiniaceae photosymbionts (Raina *et al*. 2013). Consistent with roles in stress responses, DMSP production is higher in heat-stressed *Acropora* (Raina *et al*. 2013) but acts to attract the coral pathogen *Vibrio coralliilyticus* to stressed corals (Garren *et al*. 2014). DMSP lyase catalyses catabolism of DMSP to the volatile compound DMS, which is not a chemoattractant for *V. coralliilyticus*. The genomes of *Acropora* spp. encode multiple DMSP lyases (Shinzato *et al*. 2021), several of which, including the miR-2022 target XM_015924586.1, show strong upregulation at 2-4h post-induction (Figure 1). The higher level of miR-2022 expression observed during larval development and its apparent down-regulation post-settlement (Figure 4) are consistent with the idea that this miRNA regulates XM_015924586.1 expression.

Given the caveat outlined above concerning differences in timing and nature of settlement cue between the present and previous (Brunner *et al*. 2026) studies, one potentially interesting general observation in the present study was that use of CCA chips to induce settlement of competent larvae immediately (within 10 minutes) resulted in major changes in the miRNA profile (Figure 4), preceding the time when changes occur in the expression profiles of some key marker genes (for example, homeobrain and Pumilio2; Figure 1). Presumably miRNAs play important regulatory roles in the major transcriptional transitions that occur during early polyp development. Several predicted miRNA-mRNA interactions are consequences of the cue-triggered immediate changes in the miRNA profile. Notably, miRNAs previously studied in *N. vectensis* (Moran *et al*. 2014) – miR-2023 and miR-2025 – were highly expressed during settlement in *A. digitifera*.

### Conservation of function between Actinaria and Scleractinia?

The recognition of conserved functional interactions is compromised when orthologous targets cannot be unequivocally recognised and these challenges become more acute with increasing evolutionary distance. The Scleractinia (hard corals) and Actiniaria (sea anemones) diverged >500mya (e.g. (McFadden *et al*. 2021) and possibly much earlier (Quattrini *et al*. 2018), so the extent to which miRNA function has been conserved between these two groups is unclear, but assumed to be very low. Despite these constraints, Praher et al. (2021) identified two putative cases of miR-based regulatory interactions being conserved between *N. vectensis* and corals; that of miR-2025 with Six3/6 and that of miR-2030 with NVE19315 (aka 2030T; seen only in *S. pistillata*). Whilst the predicted regulatory interaction between miR-2025 and Six3/6 identified by Praher et al. (2021) was not replicated in our analysis, this was a consequence of more stringent cutoffs being applied in the present case (see “miRNA target prediction” in Methods) – application of the original cutoff did, however, predict the proposed interaction (data not shown).

Three endogenous targets of miR-2022 were identified in *N. vectensis* by Moran et al. (2014) – NVE18870 (aka NR2; EDO30643.1), NVE16448 and NVE16498 – all of which encode constituents of nematocysts. NVE18870 encodes a nematogalectin-related protein that is a structural component of nematocysts and, whilst the function of NVE16498 is unknown, NVE16448 encodes an arylsulfatase. Subsequently Fridrich *et al*. (2023) demonstrated that miR-2022 is a key regulator of nematocyst biogenesis in *N. vectensis* and proposed that this regulatory relationship is likely to be conserved across the Cnidaria, but we have been unable to find a clear *A. millepora* homolog of *N. vectensis* NR2.

In *N. vectensis*, regulation of NVE16448 by miR-2022 occurs incoherently; the miRNA and its target are simultaneously expressed in the same cells. Although the cell-type specificity of miR-2022 expression in *A. digitifera* is unknown at present, this and its arylsulfatase (NVE16448) homolog (XP_015767398.1 = XM_015911912.1) are expressed synchronously, as in *N. vectensis*. Up-regulation of the *A. digitifera* arylsulfatase was observed between 45hpf and 69hpf (Figure 1) matching the window of expression of miR-2022, with levels of both the miRNA and its target falling rapidly after induction of settlement.

### Several *A. digitifera* miRNAs may function via TRIM71/LIN41 genes

The TRIM71/LIN41 family of E3 ubiquitin protein ligases are miRNA effectors that are ubiquitous across, but restricted to, the Metazoa and have acquired critical roles as regulators of developmental transitions in diverse members of this kingdom (Maller Schulman et al. 2008; Chang et al. 2012; Ecsedi and Grosshans 2013; Worringer et al. 2014; Tocchini and Ciosk 2015; Aeschimann et al. 2019; Spike et al. 2022). For example, members of this protein family have important roles in the regulation of both the oocyte to embryo (Spike *et al*. 2022) and the juvenile to adult (Aeschimann et al. 2019) developmental transitions in *Caenorhabditis*. During the early development of *A. millepora*, upregulation of a member of this gene family (XP_029198843.2/XP_029198844.2) coincided with major transcriptional changes that parallel the juvenile to adult transition in *Caenorhabditis* (Brunner *et al*. 2026). In the present analysis, levels of several miRNAs predicted to target (table S7) TRIM71/LIN41 gene products peak at points consistent with roles in regulating major transitions. In summary, the work presented here and in Brunner *et al*. (2026) suggest that, as in several other metazoans, TRIM71/LIN41 proteins may have important regulatory functions in developmental transitions in cnidarians, justifying more extensive investigation of their roles in this context.

## Acknowledgements

The authors gratefully acknowledge the support of the Australian Research Council Centre of Excellence grant CE14100020 as well as Discovery grants DP170104734 and DP240102310. We are particularly grateful for the support and encouragement provided by Prof Mary Collins, the former Provost of the Okinawa Institute of Science and Technology (OIST). The field work was facilitated by the OIST Marine Center, and much of the laboratory work was carried out in the laboratory of Prof Noriyuki Satoh at that institution. We gratefully acknowledge the assistance of all the staff of the OIST Marine Center with coral maintenance and other general support, and sustenance provided by the “Honey, We’re Good” noodle bar in Onna Village, Okinawa.

## DATA ACCESSIBILTY AND BENEFIT-SHARING

In accordance with database (miRbase.org) requirements, miRNA sequence data for *A. digitifera* will be lodged with miRbase after acceptance for publication. Code used for recognition, filtering and manipulation of miRNA data are available at https://github.com/milag6/Acropora-digitifera-microRNA.

## AUTHOR CONTRIBUTIONS

MG – project design, field work, molecular and bioinformatic analyses,

AF – miRNA prediction and filtering

IC – bioinformatic analyses

YM – project design, writing

RH – mRNA analyses

RB, NA-R and NU – field work

DJM – project coordination, data analyses and, with EEB, manuscript preparation

## FIGURE AND TABLE LEGENDS

**Figure S1.**
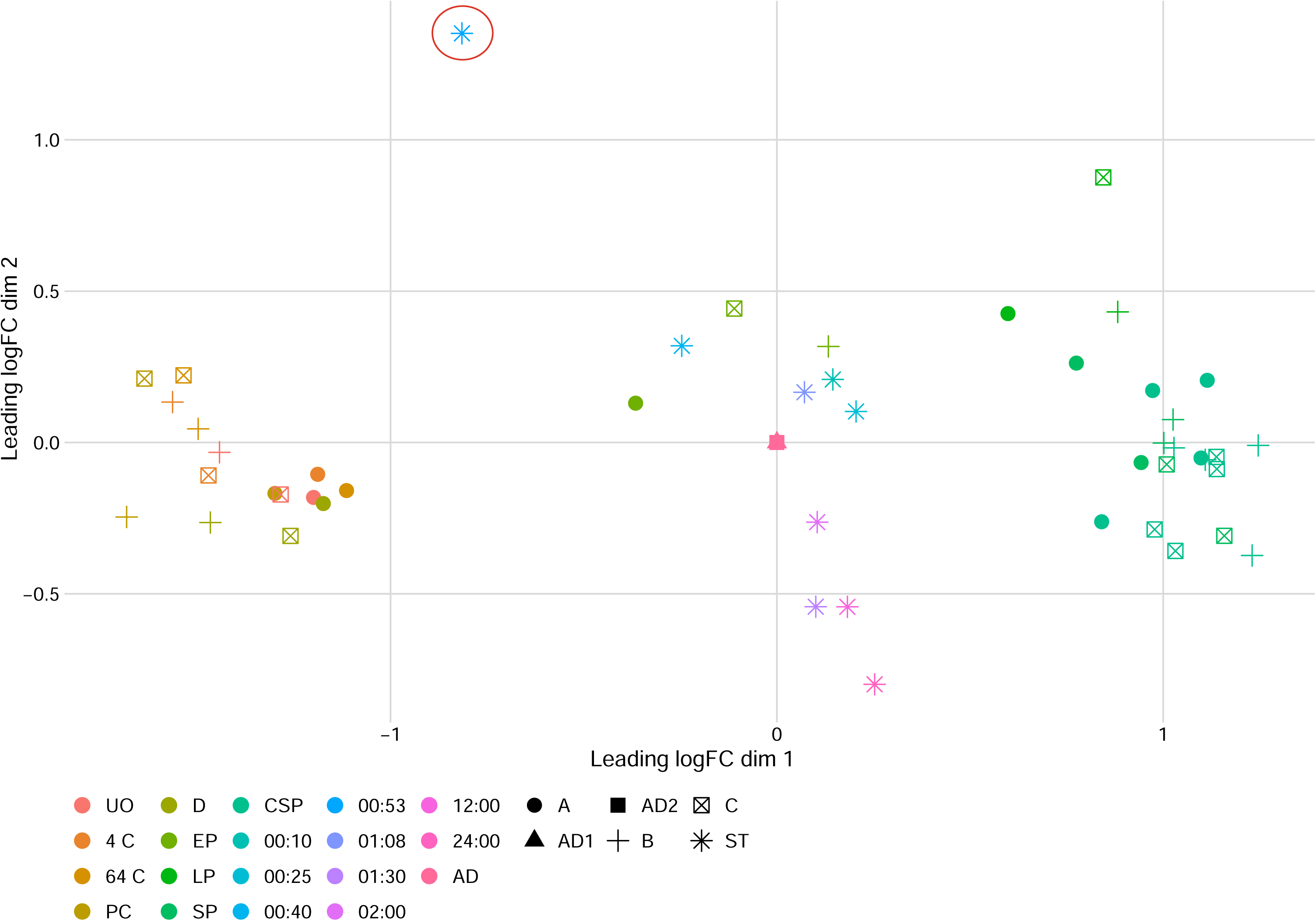
MDS ordination of miRNA expression including the settlement sample **AD_45** (00:53 after introduction of the settlement cue, CCA) that was excluded from downstream miRNA analyses. A-C labels indicate sample provenance (tanks A–C, adult colonies AD1–AD2, and the settlement experiment ST), as shown in the key. Colours indicate developmental stage abbreviated as in Figure 3 or, for post-settlement samples, time after introduction of CCA chips (hh:mm).

**Figure S2.**
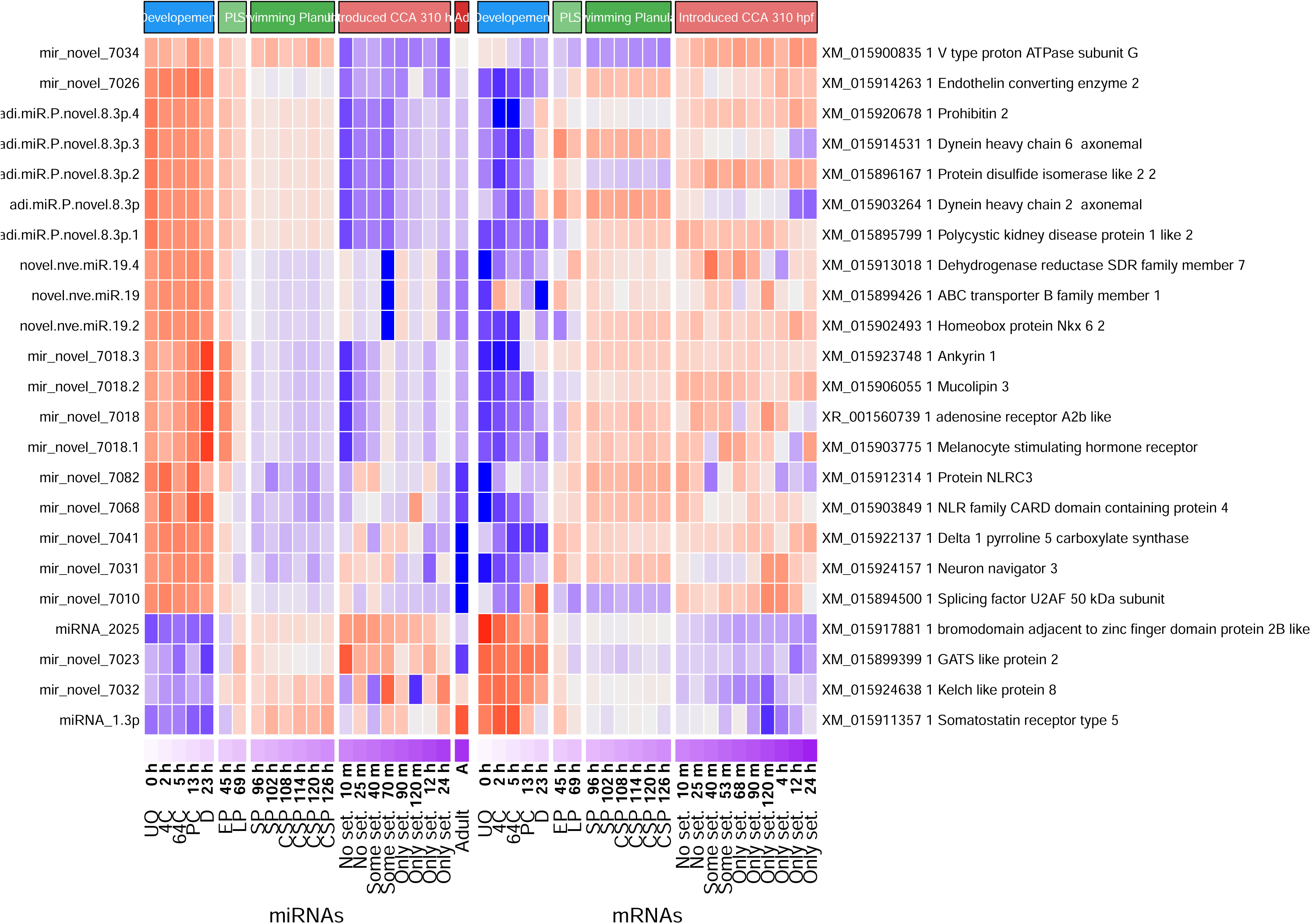
Heatmaps showing expression patterns for negatively correlated miRNA–mRNA pairs (correlation coefficient ≤ −0.5) identified when correlations were calculated across the full set of stages/time points (Supplementary Table S4). The left panel shows miRNA expression whereas that on the right shows expression of the corresponding predicted target mRNAs, aligned by sample order. Only targets with functional annotation are shown.

**Figure S3.**
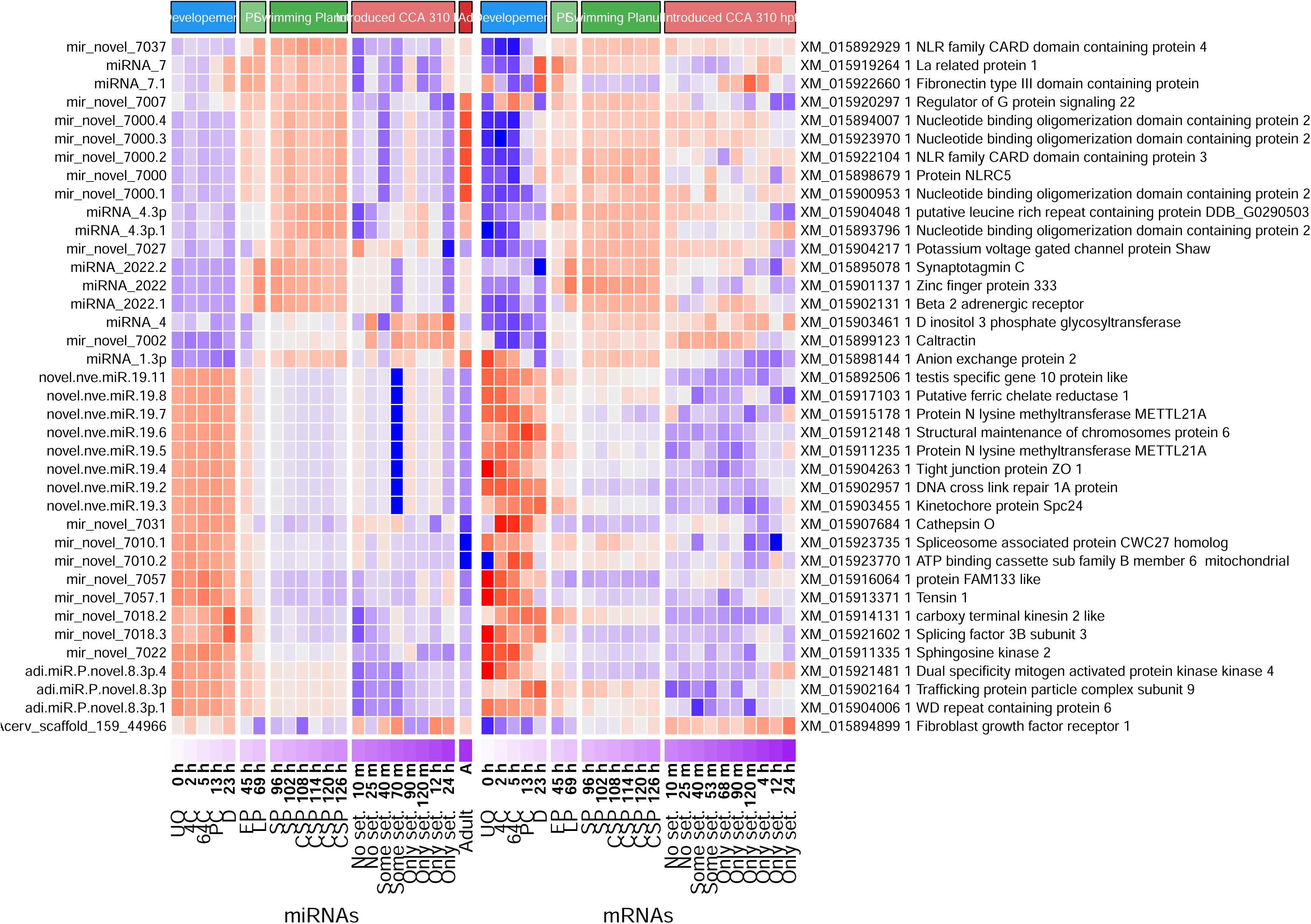
Heatmaps showing expression patterns for positively correlated miRNA–mRNA pairs (correlation coefficient ≥ 0.5) identified when correlations were calculated across the full set of stages/time points (Supplementary Table S5). The left panel shows miRNA expression and that on the right shows expression of the corresponding predicted target mRNAs, aligned by sample order. Only targets with functional annotation are shown.

## SUPPLEMENTARY TABLE LEGENDS

**Table S1.** Summary of data for miRNA-seq libraries and mapping statistics. The table lists each sample and associated metadata (including stage/time point, tank, and settlement status), with time recorded as hours post fertilisation (hpf) for pre-settlement samples and as time after introduction of the settlement cue (CCA chips) for post-settlement samples. Sequencing and filtering fields include PF (pass filter) clusters, PF reads, percent of bases passing Q30 score (%Q30), trimmed read counts, and collapse statistics, together with read mapping categories from the miRNA mapping pipeline (mapped versus unmapped).

**Table S2.** Summary of data for mRNA-seq libraries and sequencing statistics. The table lists each sample and associated metadata (including stage/time point and tank), with time recorded as hours post fertilisation (hpf) for pre-settlement samples and as time after introduction of the settlement cue (CCA chips) for post-settlement samples. Sequencing and filtering fields include PF (pass filter) clusters, PF reads, percent of bases passing Q30 score (%Q30), bases and reads after filtering, GC content, read length, and duplication level.

**Table S3.** Compiled reference list of previously reported cnidarian miRNAs used as the ‘known miRNA’ set. Entries include miRNA name, mature sequence, source species, and associated publication metadata (authors/year, journal, title, and DOI where provided).

**Table S4.** Differential expression status of predicted miRNAs relative to the low-read threshold used for downstream analyses. Each miRNA is labelled as differentially expressed or filtered out.

**Table S5.** List of miRNA loci predicted by miRDeep2 for *A. digitifera*. The table includes mature and star sequences, precursor genomic coordinates, and miRDeep2 prediction metrics (including score, read counts, and estimated probability). Naming fields indicate whether each prediction matches a previously described miRNA (retaining the original miRNA number, with species initials added where needed) or is novel (assigned a new identifier).

**Table S6.** Identities of *Acropora digitifera* genes used in comparative developmental analyses. The table lists database identifiers of those miRNA processing genes and developmentally significant genes used in the comparative analysis with *A. millepora*. In most cases reciprocal Blast analyses confirmed orthology, however, in a few cases (e.g. TRIM71/LIN41), strict orthology could not be confirmed.

**Table S7.** Predicted miRNA targets with functional annotation. This table records miRNA–target pairs from the target prediction output (including inhibition type, expectation score, target site coordinates, and the alignment/fragment sequences) and adds annotation fields for the target transcript/protein (including UniProt identifiers and protein/gene name fields where available). Red background and text indicates those miRNAs conserved across all five cnidarian species (three *Acropora* spp., *Stylophora pistillata* and *Nematostella vectensis*) and green background/text indicates novel miRNAs.

**Table S8.** Negatively correlated miRNA–mRNA pairs used for the main correlation heatmap (Figure 5). For each miRNA–target pair, the table provides the correlation statistic (stat; values ≤ −0.5), its p-value, and target annotation fields (including UniProt and protein/gene naming fields, where available).

**Table S9.** Positively correlated miRNA–mRNA pairs used for the main correlation heatmap (Figure 6). For each miRNA–target pair, the table provides the correlation statistic (stat; values ≥ 0.5), its p-value, and target annotation fields (including UniProt and protein/gene naming fields, where available).

**Table S10.** Negatively correlated miRNA–mRNA pairs identified when correlations were calculated across the full set of stages/time points (Figure S2). The table provides the correlation statistic (stat; values ≤ −0.5), p-value, and target annotation fields.

**Table S11.** Positively correlated miRNA–mRNA pairs identified when correlations were calculated across the full set of stages/time points (Figure S3). The table provides the correlation statistic (stat; values ≥ 0.5), p-value, and target annotation fields.

